# Structural and functional brain changes following four weeks of unimanual motor training: evidence from behaviour, neural stimulation, cortical thickness and functional MRI

**DOI:** 10.1101/088302

**Authors:** Martin V. Sale, Lee B. Reid, Luca Cocchi, Alex M. Pagnozzi, Stephen E. Rose, Jason B. Mattingley

## Abstract

Although different aspects of neuroplasticity can be quantified with behavioural probes, brain stimulation, and brain imaging assessments, no study to date has combined all these approaches into one comprehensive assessment of brain plasticity. Here, 24 healthy right-handed participants practised a sequence of finger-thumb opposition movements for 10 minutes each day with their left hand. After four weeks, performance for the practised sequence improved significantly (p < 0.05 FWE) relative to a matched control sequence, with both the left (mean increase: 53.0% practised, 6.5% control) and right (21.0%; 15.8%) hands. Training also induced significant (cluster p-FWE < 0.001) reductions in functional MRI activation for execution of the learned sequence, relative to the control sequence. These changes were observed as clusters in the premotor and supplementary motor cortices (right hemisphere, 301 voxel cluster; left hemisphere 700 voxel cluster), as well as sensorimotor cortices and superior parietal lobules (right hemisphere 864 voxel cluster; left hemisphere, 1947 voxel cluster). Transcranial magnetic stimulation over the right (‘trained’) primary motor cortex yielded a 58.6% mean increase in a measure of motor evoked potential amplitude, as recorded at the left abductor pollicis brevis muscle. Cortical thickness analyses based on structural MRI suggested changes in the right precentral gyrus, right post central gyrus, right dorsolateral prefrontal cortex and potentially the right supplementary motor area. Such findings are consistent with LTP-like neuroplastic changes in areas that were already responsible for finger sequence execution, rather than improved recruitment of previously non-utilised tissue.

## Introduction

The brain is capable of remarkable structural and functional change to allow it to optimise performance. Such changes, collectively referred to as plasticity, are key to understanding a variety of real-life phenomena, such as the learning of new skills [Sanes and Donoghue, 2000], the storing of memories [Xu et al., 2009], and recovering neurological function after brain injury [Nudo et al., 1996].

By definition, plasticity results in functional and/or structural change [Pascual-Leone et al., 2005]. At the microscopic level, plasticity in the central nervous system can manifest in several different ways. Sprouting of new connections, unmasking of hidden or inhibited synaptic connections, and withdrawal of inhibition are some examples of these changes [Barron et al., 2016; Hoy et al., 1985; Kong et al., 2016]. Another form of plasticity is referred to as long-term potentiation (LTP), which reflects an increase in synaptic efficacy [Bliss and Collingridge, 1993; Matsuzaki et al., 2004]. Behaviourally, neuroplastic changes typically manifest as altered task performance, such as improved speed of movement. Measuring performance of learned tasks before and after training provides a means of assessing the functional impact of training. Although such an approach can allow for an investigation of the effects of various factors on plasticity (e.g. aging, attention, hormones), a purely behavioural approach is unable to shed light on the location and type of brain changes that occur after training. In order to probe the biological changes subserving plasticity, brain stimulation and brain imaging techniques are required, each with their own relative advantages and disadvantages [Reid et al., 2016].

In the area of motor-skill learning, almost all studies investigating neuroplasticity have, so far, relied on animal models, cross-sectional data, or functional measures of brain change [Chang, 2014]. In rodents, motor learning appears to upregulate synaptic plasticity in the primary of motor cortex [Rioult-Pedotti et al., 2000], and may also be associated with altered neural recruitment in this area [Costa et al., 2004]. In humans, early work using structural magnetic resonance imaging (MRI) suggested that differences in morphology of the primary motor cortex may exist between expert musicians and the general population [Amunts et al., 1997]. Since then, functional MRI (fMRI) studies have reported altered activity in motor planning in professional sports people [Milton et al., 2007], and transcranial magnetic stimulation (TMS) studies have similarly reported that skilled racquet players have a larger cortical representation of the hand than the general population [Pearce et al., 2000]. Functional MRI work has also suggested that the supplementary motor area, dorsolateral prefrontal cortex (dlPFC), and caudate nuclei may play integrated roles in error correction and learning of motor sequences [Chevrier et al., 2007; Kübler et al., 2006].

Longitudinal studies are particularly useful for studying neuroplasticity as they eliminate the possibility that neural differences are the cause, rather than result, of participants deciding to learn a skill. Several reports exist of changes in brain function in response to motor training [Chang, 2014], including in the pre-and post-central gyri, dlPFC and basal ganglia [Floyer-Lea and Matthews, 2005]. A small number of reports have detailed minute structural changes in grey matter in response to visuo-motor tasks, particularly in the sensorimotor cortex [Bezzola et al., 2011], visual cortex, and superior parietal lobule [Draganski et al., 2004; Scholz et al., 2009], all areas associated with visuo-motor skills.

So far, research in this area has predominantly relied on single data collection modalities, such as TMS, or fMRI. Unfortunately, the differences in study design, contrasts, time-period, and imaging modalities of these studies can make it difficult to achieve a deeper understanding of the critical experimental parameters that yield reliable brain changes, whether such changes are typically concurrent or independent, and so on. In addition, each modality has its own limitations regarding reliability or interpretability, which can hamper insight into biological processes when only a single modality is acquired. For example, due to its indirect method of measuring brain activity, it is difficult to unambiguously interpret changes in fMRI in terms of biology without independent concurrent information, particularly given that controlling for task equivalency across scans can be difficult to achieve [Reid et al., 2016]. Collecting multiple modalities can also alleviate concerns that results in one modality simply reflect statistical anomalies, rather than subtle brain changes. For these reasons, the longitudinal study of motor learning presented here concurrently acquired data on four different measures of neuroplasticity – behaviour, TMS, fMRI, and cortical thickness. Findings from a fifth measure, diffusion MRI, appear in a second study [Reid et al.,]. Each of these modalities helps to build a global picture of the changes that occur with motor training, by reporting on a slightly different aspect of the neuroplastic response.

TMS can evoke a clearly discernible and quantifiable motor response when motor cortical neurons are stimulated at sufficient intensity [Hallett, 2000]. This TMS-evoked response, referred to as a motor evoked potential (MEP), reflects the excitability of underlying cortical neurons. Measuring the amplitude of the MEP before and after a training paradigm provides objective measurement of changes attributed to cortical motor plasticity [Stefan et al., 2000]. This is different from other, non-motor regions, where brain stimulation-evoked responses are more difficult to quantify (e.g., prefrontal brain regions), yet which also undergo changes via the same, ubiquitous, mechanisms. Although TMS is ideally suited to quantify training-related change arising in the motor cortex, there are several limitations to the technique. In particular, TMS can only penetrate superficial grey matter structures [Roth et al., 2007], and can only probe the activity of one area of cortex at a time. Therefore, TMS is unable to provide information on whole-brain changes arising from training.

Whole-brain assessment of plastic change arising with training can be investigated with both functional and structural MRI. Functional MRI of local blood oxygenation level-dependent (BOLD) signals provide information about changes in brain activity during tasks with high sensitivity and excellent spatial resolution. However, fMRI cannot readily distinguish whether any measured changes following training reflect excitatory or inhibitory activity [Logothetis, 2008], something that TMS can do. Changes in BOLD signals are also more straightforward to interpret, in terms of biological change, when in the context of information from other modalities, such as TMS [Reid et al., 2016]. By contrast, if conducted carefully, structural MRI can measure changes in cortical thickness that are more directly interpretable in terms of biology but, without accompanying functional measures, are difficult to link to any changes in brain activity.

With these points in mind, the aim of the present study was to use a multi-modal approach to investigate plasticity in the human cortex. We measured behaviour, TMS evoked MEPs, BOLD changes using fMRI, and cortical thickness, before and after four-weeks of motor training. To make interpretation of the results more straightforward, we opted for training that did not have any visual component. In combination with our parallel analysis of white matter and network changes [Reid et al.,], this work provides a comprehensive assessment of the functional and structural brain changes associated with motor learning in humans.

## Materials and Methods

### Overview

Twenty-four participants were recruited (14 female; aged 28.8 ± 1.5 years; range 18 – 40 years). Participants were all right-handed (laterality quotient 0.92 ± 0.03; range 0.58 – 1.0) as assessed by the Edinburgh Handedness questionnaire. Participants trained daily on a finger-thumb opposition task [Karni et al., 1995] for four weeks. Behavioural, TMS, and MRI measures were obtained before and immediately after the training period to quantify training-related changes. No participants reported any adverse effects. The study was approved by the University of Queensland Human Research Ethics Committee and all participants gave written informed consent.

### Motor training task

The training task has been used in a previous independent study, in which it was reported to induce robust behavioural and functional changes [Karni et al., 1995]. The task involved participants performing a sequence of finger-thumb opposition movements with their non-dominant (left) hand (Figure 1). Participants were pseudo-randomly assigned one of two sequences that were mirror-reverse copies of each other. Participants were instructed to perform their assigned sequence as quickly and as accurately as possible for 10 minutes each day for 4 weeks, and not to look at their hand during training. To help remember the correct sequence, participants were given a printed copy of their allotted sequence (red or blue; see Figure 1). In order to minimize any circadian effects on motor learning [Sale et al., 2008], each participant was randomised to practising either during the morning or evening, and conducted training at the same time each day. Participants kept a log-book to record their training, and were instructed to be honest when training sessions were missed, or were performed outside of the specified training time.

**Figure 1.**
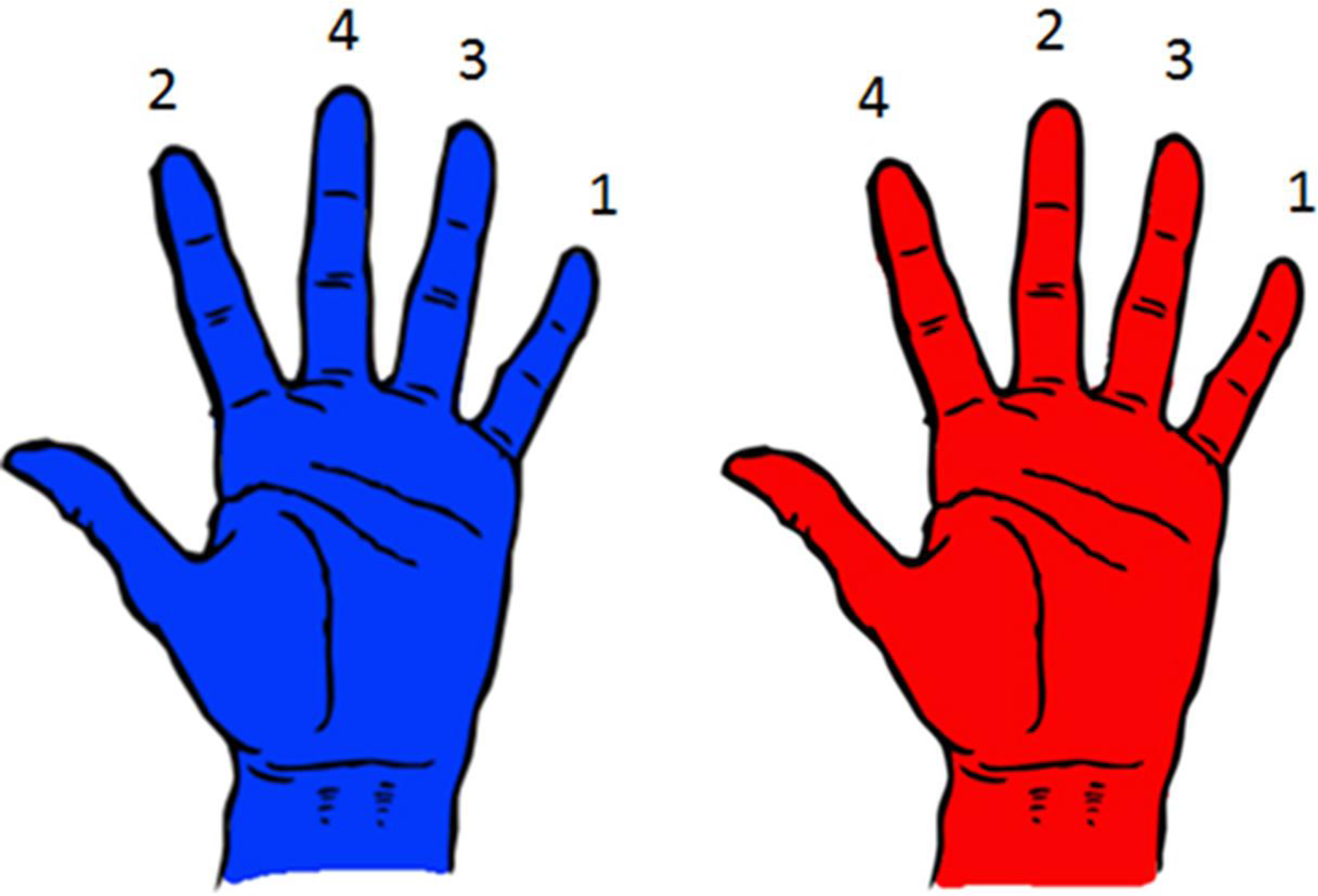
Participants were randomised to practising one of two finger-to-thumb opposition sequences. For the ‘Blue’ sequence (left), the order of fingers required to make contact with the thumb were little, index, ring, middle. For the ‘Red’ sequence (right) the order was little, middle, ring, index. These sequences were mirror images of one another.

### Behavioural measure

Participants’ performance on the finger-thumb opposition tasks was assessed as the number of correct sequences completed in 30 sec. Performance was documented online with a hand-held video camera, and quantified off-line. Participants performed both red and blue (Figure 1) sequences with both their left and their right hands. This was to investigate whether training induced any spill-over of effects to a non-trained sequence and/or the contralateral hemisphere. The order of the sequences performed was randomised for each participant. To minimize errors, and to assist in performance, a print-out of the sequence to be performed was placed in front of the participant for the duration of that task.

Performance on the finger-thumb opposition sequences was analysed using a three-way repeated measures ANOVA with within-participant factors of *training* (Baseline, Post), *hand* (Left, Right), and *sequence* (Trained, Control). Where appropriate, post hoc analyses were performed using Holm-Bonferroni corrected paired t-tests.

### Cortical Thickness

We examined cortical thickness changes arising in response to motor training. We acquired MPRAGE T1 images (0.9mm isotropic) immediately before (“baseline”) and after (“post”) the four-week training period using a 3T MR system (Magnetom Trio, Siemens) and a standard 32 -channel head coil. Images were processed with Advanced Normalization Tools (ANTs; v2.1.0, source pulled 9th Feb 2016). An overview of the processing pipeline is shown in Figure 2.

**Figure 2.**
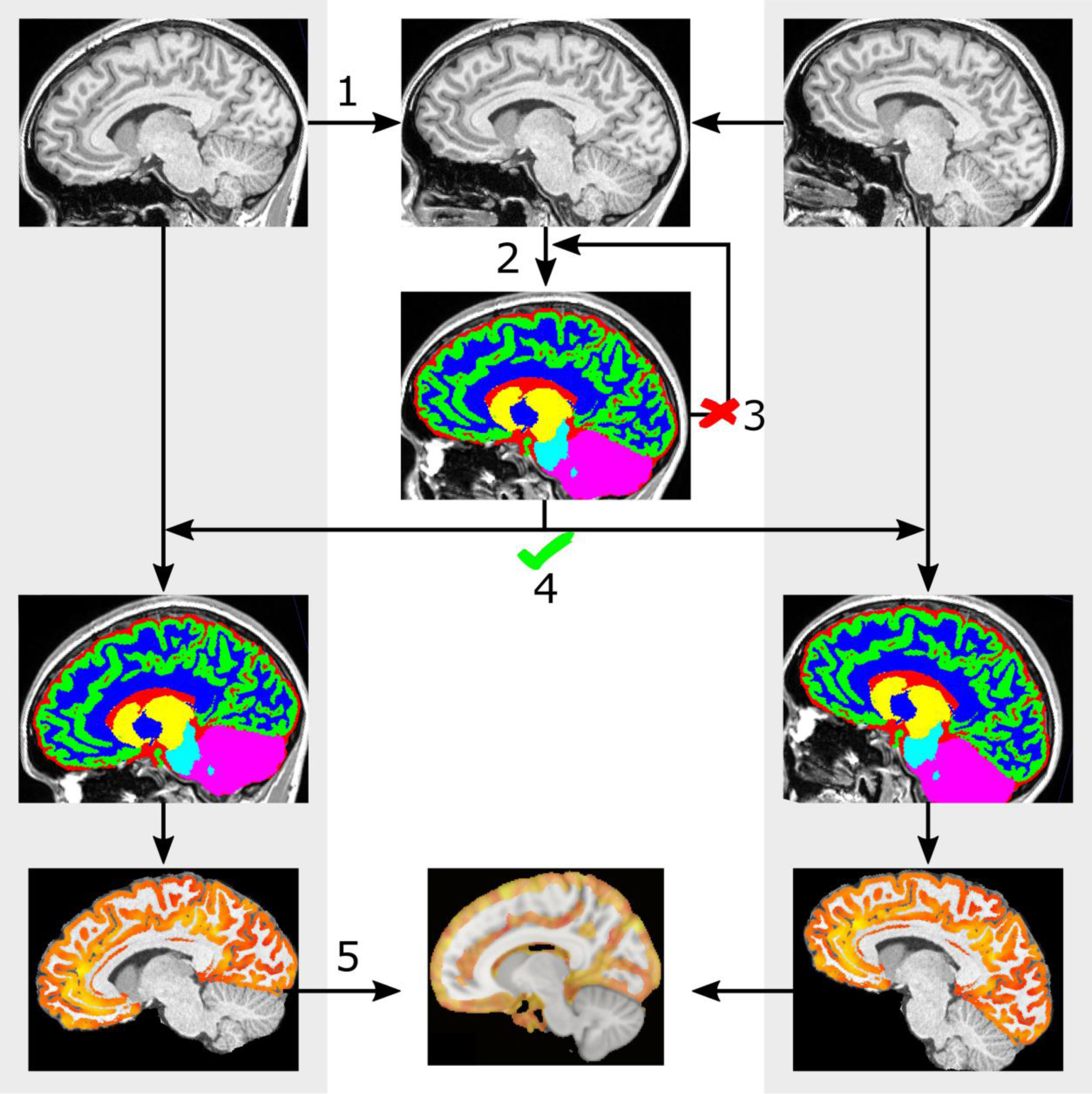
Cortical thickness pipeline. Steps are indicated with numbers. 1) Structural volumes from the baseline (left column) and post (right column) were pre-processed, skull-stripped, and registered using a symmetrical registration. The resulting half transforms were applied to non-skull-stripped images to achieve an intermediate image (top, middle column), which was processed into a sharp single subject template. 2) The single-subject template was then skull-stripped and segmented. The result was carefully visually inspected. 3) If the segmentation was considered inaccurate, the brain mask was manually edited and Step 2 was re-run. 4) If the segmentation was successful, each time point was then re-skull-stripped and segmented using the single subject template (middle row of left and right columns), and cortical thickness calculated (bottom row of left and right columns). 5) Cortical thickness at each time point was moved into single-subject template space, subtracted, and this difference transformed into MNI space for statistical analysis using the known transform between the single-subject template and MNI space. Statistical analyses were then performed across all subjects, resulting in statistical maps (bottom row, middle column). Images here are purely illustrative; orientation differences have been exaggerated in order to convey concepts clearly.

#### Single Participant Analyses

In longitudinal analyses, it is important to utilise templates that are invariant to either time-point, in order to boost statistical power and avoid false-positives induced by registration bias [Thomas et al., 2009]. Although ANTs provides a longitudinal cortical thickness script, we found that the templates produced by this script on our system were commonly biased toward one time-point. Therefore, we constructed templates for each participants by rigid-registering the baseline and post structural images together using ANTS SyN (symmetric) transform [Avants et al., 2008] after skull-stripping, N4 bias-field correction and intensity normalisation. Time points were transformed by half of the resulting matrices then averaged, producing an approximate template that was unbiased with respect to time point. This initial template was converted into a sharper, segmented, template using the *antsMultivariateTemplateConstruction* script, followed by the *antsCorticalThickness* script using the ANTs NKI template. Each axial, coronal, and sagittal slice was then carefully visually inspected. In instances where dura or skull were classified as grey matter, or the brain under-segmented, edits were made to the brain mask using ITK Snap v3.4.0 [Yushkevich et al., 2006], and the script was re-run with the updated brain mask. For this study, particular emphasis was placed on achieving highly precise segmentations of the parietal and frontal lobes (Supplementary Figure 1) due to their known association with, or proximity to, areas involved in motor training and/or learning. The process was repeated until we were satisfied with the final segmentations. Examples of acceptable and unacceptable segmentations are provided in Supplementary Figure 2. Posterior tissue probabilities were then converted into priors in a manner consistent with ANTs *antsCookTemplatePriorsCommand* script.

The *antsCorticalThickness* script was applied to structural images from both time points, utilising the single subject template, producing a cortical thickness map in single-subject-template space. Subtraction of the pre-training from post-training cortical thickness images produced a cortical thickness difference image.

#### Statistical Analysis

We hypothesised that any region of change would likely be substantially smaller than atlas ROIs, and so opted for voxel-based morphometry. Single subject template T1s were registered to FSL’s 1mm isotropic MNI152 atlas (Figure 2) using ANTs SyN, and the resultant transforms applied to cortical thickness difference images. Voxels where the white matter was the most probable tissue, as defined by the mean of all single-subject tissue priors, were excluded. Images were then smoothed with a 5mm FWHM kernel and placed into a factorial model in SPM 12 (http://www.fil.ion.ucl.ac.uk/spm/software/spm12/). This model regressed change in cortical thickness against time point. This model included sex as a factor due to account for its previously-reported effects on cortical thickness [Luders et al., 2006], and included ANCOVA normalisation for nuisance effects to account for any remaining global differences. In the interests of accuracy and statistical power, we restricted our analysis to the parietal and frontal lobes – the areas which received particular focus during segmentation correction (Figure 2). We set our significance criteria as a p<0.05 FWE corrected, or a cluster comprising more than 20 voxels expressing values p<0.001 uncorrected.

### Functional MRI

Functional MRI images were acquired in the same session as structural MRI images. We acquired 41 axial slices (slice thickness, 3.3 mm) using a gradient EPI sequence (TR 2.67 sec; TE, 28 ms; flip angle, 90°; field of view, 210 x 210 mm; voxel size, 3.3 x 3.3 x 3.3 mm). A liquid crystal display projector back-projected the stimuli onto a screen positioned at the head-end of the scanner bore. Participants lay supine within the bore of the magnet and viewed the stimuli via a mirror that reflected the images displayed on the screen. Participant head movement was minimized by packing foam padding around the head.

Prior to entering the scanner, participants familiarised themselves with the two movement sequences – ‘trained’ and ‘control’ – they were going to perform within the scanner. The sequences are shown in Figure 1. Within the scanner, participants performed these finger-thumb opposition movements with their left hand in blocks of 16 seconds, each followed by 16 seconds of rest. During rest blocks, a visual display showed a “Rest” command. At the start of each movement block a visual cue - displaying either the red or blue hand and corresponding movement sequence - notified participants whether they would be performing the trained or the control sequence. This was displayed for 2 seconds, then removed. Participants then performed the required sequence at a rate of two movements per second. As a cue to aid in this timing, a fixation cross flashed at 2 Hz on the screen. A tone also sounded at 2 Hz intervals throughout the acquisition. The last two tones in each movement block were at progressively lower pitches to notify participants that a rest block was imminent. Immediately following completion of each movement block (i.e., following the last fixation cross), a “Stop” command was presented for 1 second, which was then replaced with the “Rest” command, Four consecutive ‘runs’ were performed. Each run consisted of four trained-sequence blocks, four control-sequence blocks, and seven rest blocks. The order of the trained/control sequence blocks was randomised but kept consistent between participants. Correct performance of the sequence was verified by recording the movements with a video camera, which were later reviewed for accuracy.

Image processing and statistical analyses were performed using SPM12 (Wellcome Department of Imaging Neuroscience, UCL, London, UK). Functional data volumes were slice-time corrected and realigned to the first volume. The mean image of the resultant time series was co-registered with the participant’s temporally-unbiased T1 template (see Methods: Cortical Thickness). The time series was then normalised into MNI space, an MNI brain mask was applied, and the result smoothed with an 8-mm FWHM isotropic Gaussian kernel. For first level statistical analyses, contrasts were conducted for (a) movement > rest (b) trained > rest, and (c) control > trained sequence blocks. All six motion parameters for each run were included as nuisance regressors. The second level analysis looked at the interaction between sequence and time point (i.e., baseline [control > trained] versus post [control > trained]). This was designed to show how the brain responded differently to the learned and control sequences after training, taking into account any differences that may have existed before training began (although no baseline differences were expected). It was possible that the trained or control sequence would show greater BOLD responses after training, and so we treated this as a two-tailed test. To account for this, we set a more conservative cluster significance level of p<0.025 FWE with minimum cluster size of 200 voxels, after voxel-wise thresholding at p<0.0005 uncorrected.

### TMS mapping

Mapping of motor cortical excitability with TMS was performed in a separate laboratory, after MRI acquisition. Surface electromyographic recordings were obtained from the left abductor pollicis brevis (APB) muscle. Recordings were made using silver/silver-chloride surface electrodes with the active electrode placed over the muscle belly, and the reference electrode placed over the adjacent metacarpophalangeal joint. Signals were amplified (x1000) and band-pass filtered (20-1000 Hz) using a NeuroLog system (Digitimer), then digitized (2000 Hz) with a data acquisition interface (BNC-2110; National Instruments) and custom MATLAB software (MathWorks). Signals were monitored online for movement-related activity using high-gain electromyography and a digital oscilloscope.

Mapping of motor cortical excitability before and after training involved applying TMS to the right hemisphere in a grid like pattern (described below). The TMS was delivered using a Magstim 200^2^ stimulator (Magstim, UK) and a figure-of-eight coil (70 mm diameter). The coil was positioned with the handle pointing backwards at a 45° angle to the sagittal plane to preferentially induce current in a posterior-to-anterior direction in the cortex. The optimal scalp position for evoking electromyographic responses in the APB was established as the position that consistently evoked the largest MEP amplitude in this muscle with a slightly suprathreshold intensity. Resting motor threshold (RMT) was determined and defined as the minimum stimulus intensity which evoked an MEP of at least 50 μV in at least 5 out of 10 successive stimuli. The TMS intensity used for the mapping procedure was set at 120% of RMT. This stimulus intensity was established for both pre-and post-training sessions.

Prior to participants’ first TMS mapping session, their individual T1 MRI scan was processed with the neuro-navigation software ASA-Lab (ANT, The Netherlands). Markers were placed on the scalp surface of the structural scan in a grid-like pattern spanning the entire right hemisphere. The grid commenced at the vertex, and markers were placed every 1 cm anteriorly and posteriorly from the vertex, and then extended out laterally (to the right) in 1 cm increments. Marker sites were targeted with TMS using a Polaris-based infrared frameless stereotaxic system and Visor software (ANT, The Netherlands). Five TMS pulses were applied to these markers every 5 sec. The markers were stimulated systematically, moving in a medial-to-lateral direction. Stimulation of the marker sites continued until the average MEP amplitude from a marker site fell below 50 μV. Those sites where average MEPs met or exceeded this amplitude were referred to as *active*. Once a row of markers had been assessed, stimulation was moved either 1cm anteriorly or posteriorly (chosen randomly), until all active marker sites had been stimulated and identified. The MEP *volume* [Schabrun et al., 2009] was also calculated before and after training. This was calculated by summing the mean MEP amplitude of all the active sites.

Training-related increases in the number of active sites and MEP volume were investigated using separate one-tailed, paired t-tests to compare pre-training measurements with those acquired after training. Significance was set as p<0.05 after Holm-Bonferroni multiple comparisons correction.

### TMS and MRI overlay

In order to interpret TMS, fMRI, and cortical thickness results together, it was important to show reasonable anatomical correspondence between the methods. To achieve this, TMS MEP responses were projected into MNI 152 space. This was achieved by converting each active site from Talairach to MNI coordinates, using the Lancaster transform [Lancaster et al., 2007], then connecting neighbouring measurement nodes to form a mesh. An edge of zero-value nodes was added to the outside of this mesh, based on the position of neighbouring nodes. A duplicate of this mesh was projected 20mm toward the midline of the base of the brain (x=0, y=0), reflecting the approximate penetration of the TMS pulse. Values on the grid were normalised between 0-1 on a per-participant basis. For each participant, a volume was generated by linearly interpolating all voxels in MNI space between the inner and outer surfaces of the grid by the nearest surrounding 8 nodes. All participants’ volumes were averaged to produce a mean image. This overlay was used for visual comparison of the modalities only; it was not used for quantification of TMS MEPs.

## Results

### Behavioural performance

Prior to training, participants were equally proficient at performing the trained and control sequences with their left and right hands. Following training, there was a significant improvement in the number of sequences participants could complete in the 30 sec period (Figure 3). Using the left (trained) hand, the number of correct sequences completed on the trained sequence increased by 53%, from 24.0 ± 1.2 (Mean ± SEM) to 36.7 ± 2.1 (p < 0.05 FWE). There were also smaller, but significant (p < 0.05 FWE) improvements in performance of the control sequence with the left hand (6.5%), trained sequence with the right hand (21.0%) and control sequence with the right hand (15.8%). Although participants improved to a greater degree at the trained sequence than the control sequence (both hands p < 0.05 FWE), this difference between sequences was substantially greater for the left than right hand (p<0.05 FWE).

**Figure 3.**
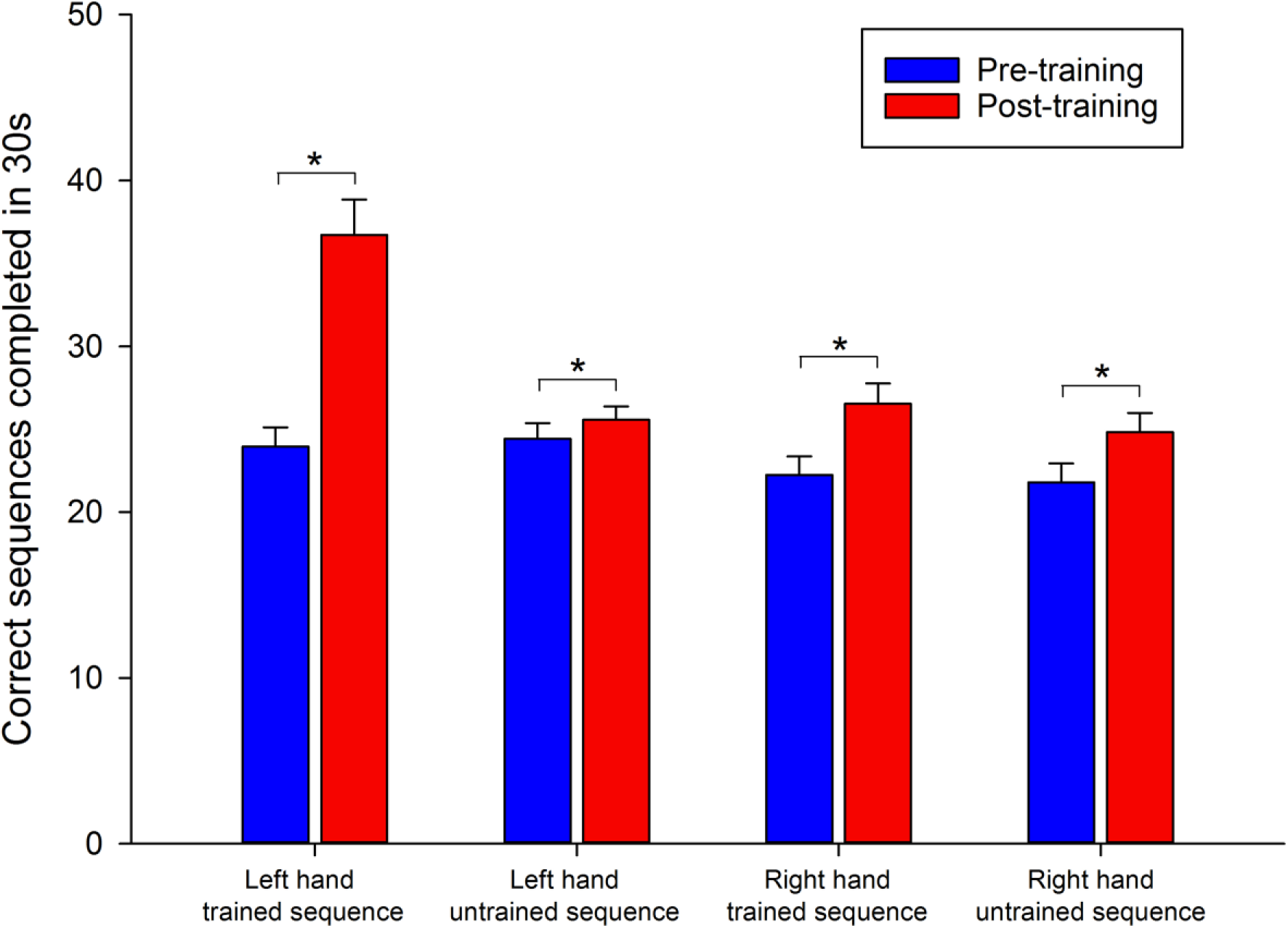
Increase in performance of motor training tasks following four-weeks of training. Group data (n = 24) showing number of correct sequences performed prior to (blue bars) and following (red bars) four weeks of training of a finger-thumb opposition movement sequence (trained sequence). Participants performed the trained and control sequence with their left hand (trained) and right hand (not trained). Training improved execution speed for all four hand-sequence combinations assessed (Each p<0.05 FWE). Data represent mean ± SEM.

### TMS mapping

There was no significant change in RMT following training (40.8% vs 41.6% maximum stimulator output; p > 0.05). By contrast, training induced an increase in the *volume* of MEP amplitudes evoked at the targeted sites by an average of 58.6% (p = 0.015; Figure 4). That is, with the same relative TMS intensity (120% of RMT), MEPs evoked by TMS at the active sites were significantly larger following training. There was no significant increase in the number of active sites evoked with TMS (p=0.09). A cortical heat-map from a representative participant is shown in Figure 4, together with group-wise changes. A normalised group-wise heat map (see Methods) is also shown in Figure 5.

**Figure 4.**
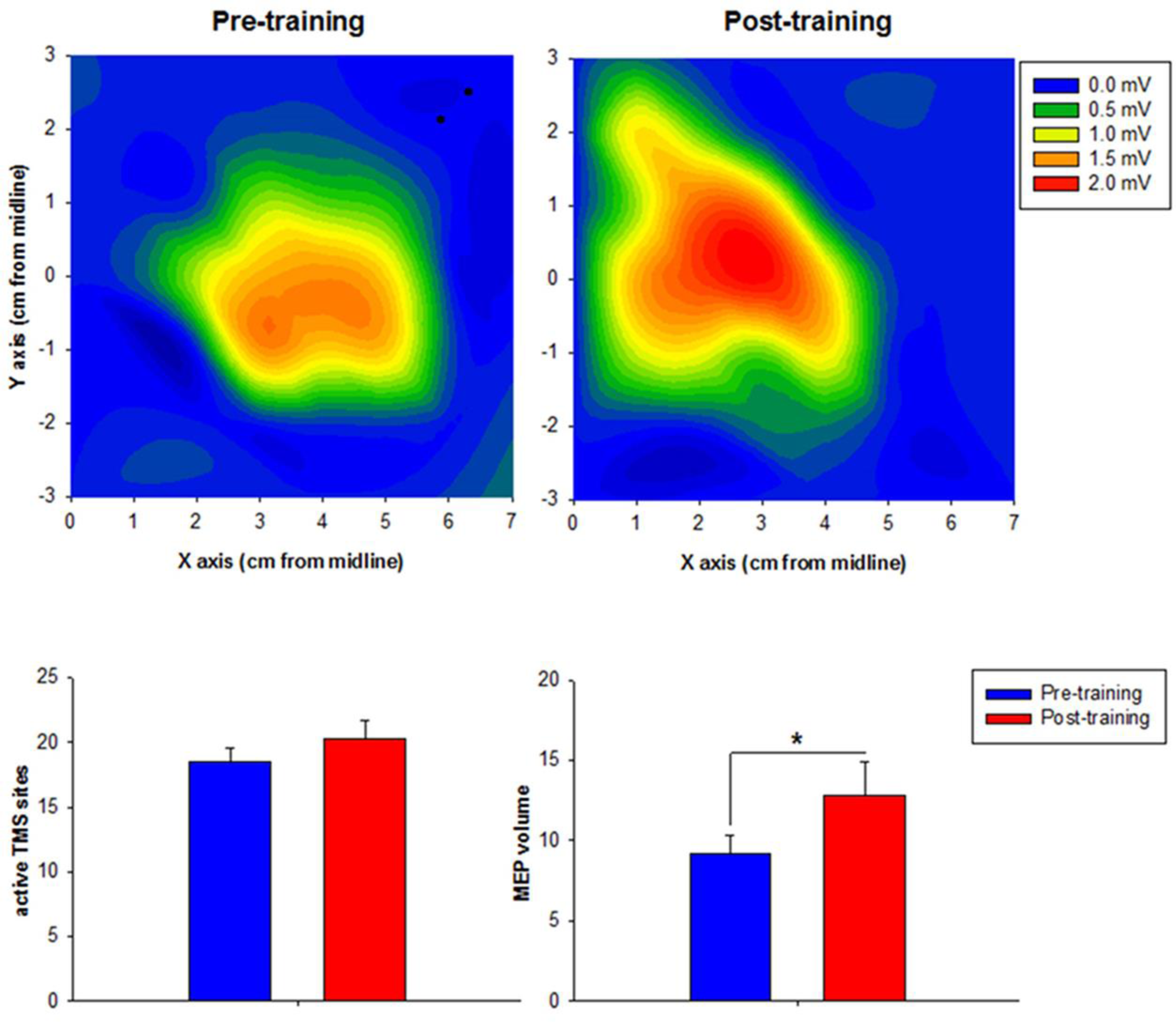
Training-related changes in motor cortical excitability. Top: A heat-map representation of the cortical representation of the abductor pollicis brevis before (left) and after (right) training in one representative participant. Coordinates are referenced to the vertex (0, 0). The average MEP amplitude evoked at each site is indicated by the colour scale (mV). Bottom: Mean number of active sites (left) and MEP map volume (right) before (blue) and after (red) training across all participants. There was no increase in active sites following training, but MEP volume increased significantly.

**Figure 5.**
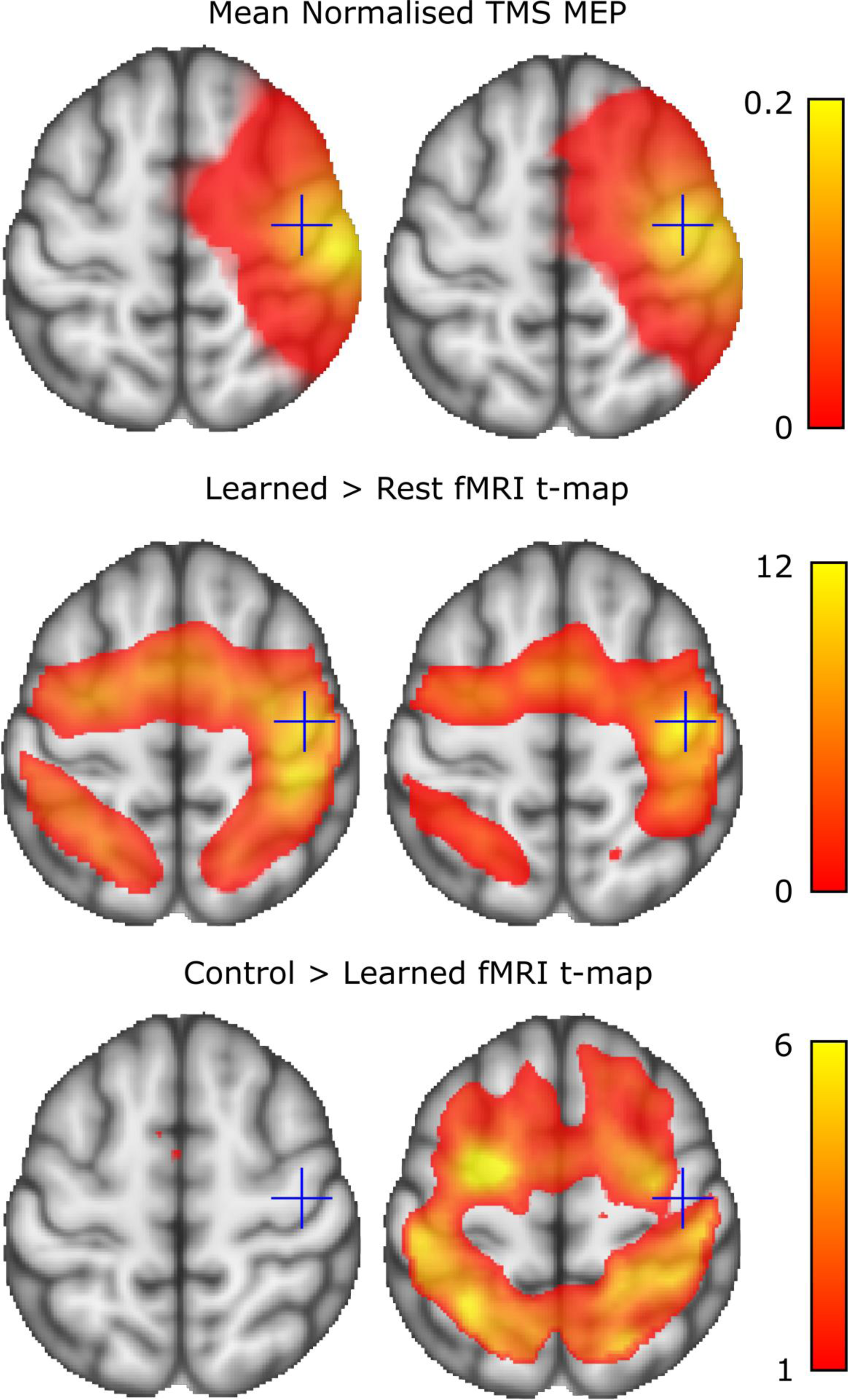
Mean of participants’ normalised TMS responses (Top; normalised mV), positive BOLD activation during the learned task (Middle; t-values for task vs rest), and positive t-value map contrasting BOLD for control sequence > trained sequence (Bottom), overlaid on an MNI 152 T1 image. The left and right columns show measures before and after the training period, respectively. Crosshairs show equivalent anatomical positions (MNI 152 coordinate: 37, -16, 57) to aid viewing. TMS and BOLD signals show a large degree of overlap in the primary motor cortex. The TMS responses were centred more anteriorly (motor+premotor), particularly after training, than the BOLD responses (predominantly motor+ sensorimotor). Details of TMS projection into MNI space is contained in the text.

### Cortical Thickness

One participant’s dataset was lost during transfer to the server, and so was excluded from both cortical thickness and fMRI analyses. A second participant displayed slight MRI artefacts on the T1 images in the right sensorimotor cortex, and so was excluded from cortical thickness analyses. For a third dataset we were unable to achieve a high-quality tissue segmentation and so we excluded this dataset from cortical thickness analyses, leaving 21 participants in total for these analyses. At an uncorrected threshold of p<0.001, training resulted in an increase in cortical thickness in the right precentral gyrus (81 voxel cluster), right post central gyrus (34 voxel cluster) and right dlPFC at approximately Brodmann’s area 9 (22 voxels; Figure 6). The right supplementary motor area (SMA) also contained two nearby clusters, 11 and 13 voxels in size, whose individual volumes did not cross our prespecified threshold, but whose sum did. Although these five clusters did not survive multiple comparisons correction (p<0.05 FWE) all changes were observed in regions that are likely to be involved with motor tasks, and all were in the right (‘trained’) hemisphere. Notably, clusters at the pre-and post-central gyri were consistent with the expected location of sensory and motor finger representations [Penfield and Boldrey, 1937; Wahnoun et al., 2015], and close to peaks in the (post time-point) TMS and fMRI maps. All clusters were within the group-wise region of successful TMS excitation, and all but the prefrontal cluster were consistent with regions of group-wise fMRI activation. To test the robustness of these findings, we removed data from the two participants who displayed the strongest performance improvements. In this reanalysis, the previously found clusters in the pre-central gyrus, post-central gyrus, and prefrontal cortex were still apparent (data not shown).

**Figure 6.**
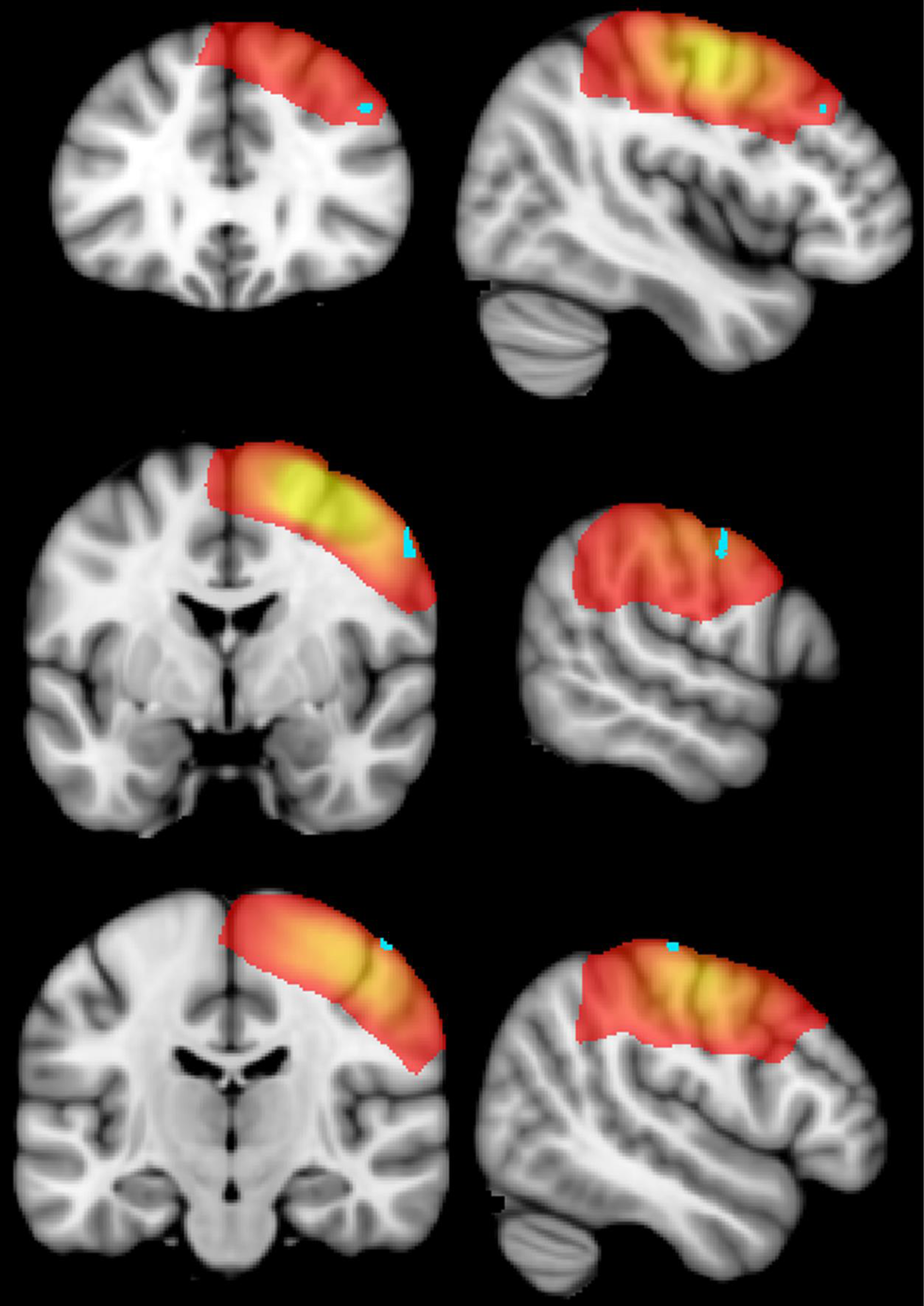
Areas of increase in cortical thickness overlaid on the FSL MNI 152 template. Light blue indicates voxels with significant (cluster size > 20 voxels at p<0.001 unc) increases in cortical thickness estimation. Increases were seen in the right prefrontal cortex (top row), and approximate hand areas of the right precentral (middle row) and postcentral (bottom row) gyri. Red-Yellow indicates mean normalised TMS response, where yellow indicates a strong response (mean ≥ 30% normalised peak motor evoked potential) and red indicates a weaker response (mean ≥ 1% of normalised peak motor evoked potential); see text for details. Left of coronal and axial images indicates left of brain. Axial slices (top-to-bottom) show MNI y-coordinates 30, -3 and -18. Sagittal slices show MNI x-coordinates 44, 58 and 61.

### fMRI analysis

Of the 23 participants included in the fMRI analysis, none exhibited excessive (>2mm or 2°) head movement during any session. At the pre-training timepoint (Figure 5), the (soon-to-be) learned sequence versus rest contrast revealed four significant (p<0.05 FWE) clusters that were located across a variety of motor areas, predominantly in a bilateral manner, including the precentral gyri, postcentral gyri, supplementary motor areas and superior parietal lobule. At this time, the activation of the (soon-to-be) trained and control sequences were, as expected, equivalent (i.e., there were no significant clusters when contrasted). Following training, however, the *control* sequence evoked *greater* activation than the trained sequence in a number of sensorimotor areas (Figure 5). To formally compare these differences we calculated the interaction between training type and time point (baseline [control > trained] versus post [control > trained]). For this contrast, positive t-values indicated locations in which the trained sequence showed reduced signal after training, relative to the control sequence. In the left hemisphere, a significant cluster overlapped the post-central gyrus and superior parietal lobule (1947 voxels, cluster p-FWE < 0.001; Table 1). In the right hemisphere, a similar cluster overlapped the pre-central gyrus, post-central gyrus, and superior parietal lobule (864 voxels, cluster p-FWE < 0.001; Figure 7). Both hemispheres also each demonstrated a significant cluster crossing the premotor cortex and supplementary motor area (left hemisphere, 700 voxels; right hemisphere, 301 voxels; each cluster p-FWE < 0.001). Reversal of this contrast, highlighting areas of increased relative activation of the learned sequence, revealed no significant clusters.

**Table 1.** Locations of significant cluster fMRI peaks for the timepoint-sequence interaction (Baseline [Control > Trained] > Post [Control > Trained]). Cluster size is in voxels. Peaks are in MNI 152 standard space. Abbreviations: L, Left; PrG, Precentral Gyrus; R, Right; SMA, Supplementary Motor Area; SPL, Superior Parietal Lobule

| Cluster Size | Peak Coordinates | Anatomical Locations |
| --- | --- | --- |
| 1947 | -26, -54, 54 | L SPL |
|  | -44, -34, 36 | L PoG |
|  | -32, -44, 52 | L SPL |
| 864 | 18, -56, 60 | R SPL |
|  | 40, -30, 42 | R PrG/PoG |
|  | 20, -48, 62 | R SPL/PoG |
| 700 | -12, -6, 64 | L SMA |
|  | -24, 6, 40 | L Premotor/SMA |
|  | -24, 10, 42 | L Premotor |
| 301 | 24, -8, 54 | R Premotor/SMA |
|  | 24, -2, 64 | R Premotor/SMA |
|  | 28, -10, 62 | R Premotor |

**Figure 7.**
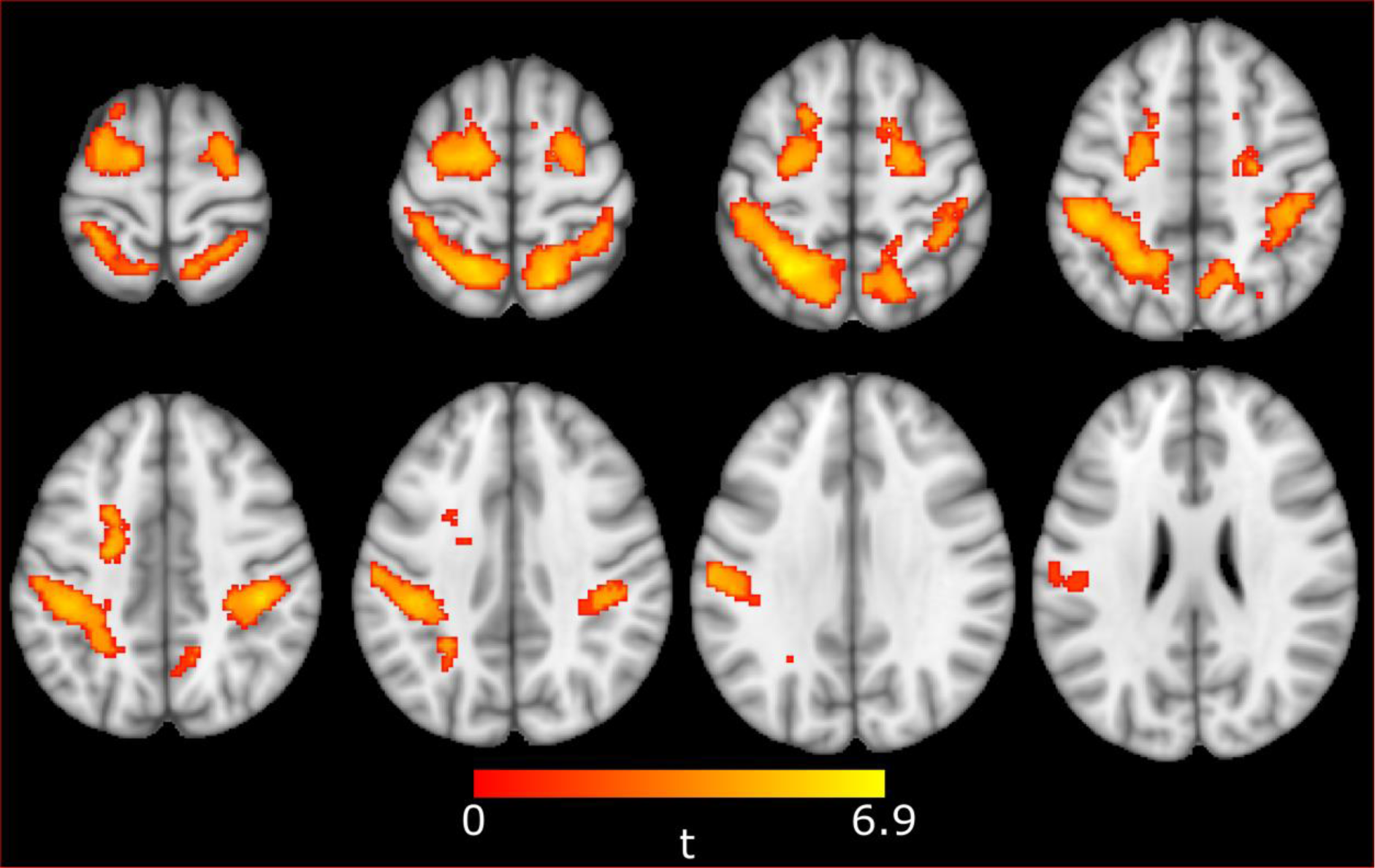
Training-related differences in BOLD activation. Groupwise functional MRI t-map, showing voxels which, after training, demonstrated higher signal during the control sequence than the trained sequence. A series of axial slices is shown. Significant clusters were apparent in the superior parietal lobule and primary sensorimotor cortices in the right (864 voxels; p<0.001 FWE) and left (1947 voxels; p<0.001 FWE) hemispheres. Significant clusters were also seen in the supplementary motor area / superior premotor area of the right (301 voxels; p<0.001 FWE) and left (700 voxels; p<0.001 FWE) hemispheres. Only voxels belonging to significant clusters are shown; a structural template is displayed for anatomical reference only. Left of image is left of brain. Axial slices shown are evenly spread between MNI-152 z-slices 67 and 26 inclusive.

## Discussion

The neural changes that accompany a period of motor training contribute to the increase in performance. Understanding how these neural changes manifest themselves is important in both guiding rehabilitation strategies [Reid et al., 2015], and understanding normal brain function. Such neural changes can be quantified in several ways, each with their respective advantages and limitations. Here, we show how utilising several different methods to concurrently quantify plasticity – behavioural, brain stimulation, functional brain imaging, and structural brain imaging – can offer broad insight into the plastic changes that arise following training. To our knowledge, these multimodal data from healthy adults provide the most comprehensive assessment of the functional and structural changes that occur following training to date.

Four weeks of training of a sequence of finger-thumb opposition movements resulted in a substantial improvement in performance, particularly of the trained sequence (Figure 3). To further investigate the brain changes arising from training, we incorporated several other probes of cortical structure and function.

### fMRI

Our fMRI analyses revealed that, only after training, the trained sequence evoked *lower* cortical activation bilaterally than the control sequence (Figure 7), predominantly in the sensorimotor cortices and superior parietal lobe (Table 1). The apparent relative reduction in functional activation reported here is at odds with an earlier smaller (n=6) study that used a very similar training and scanning approach [Karni et al., 1995]. A subsequent replication of this work showed that fMRI activation decreased during the 3^rd^ and 4^th^ weeks of training, and PET data suggested that this was, at least in part, due to an increase in baseline blood flow rather than a decrease in actual brain activity [Xiong et al., 2009]. Changes in baseline blood flow may certainly have taken place in the present study, but are unlikely to explain our results because our contrast focused on the interaction between time point and task. Specifically, since both the control and learned tasks evoked the same patterns of activation at baseline (Figure 5), changes in resting cerebral blood flow would be expected to affect both equally, and so cannot explain the statistical interaction reported here. An alternative, non-mutually exclusive, explanation is that local changes, such as LTP, allowed local grey matter to perform the learned sequence more efficiently, reducing the need for recruitment of surrounding areas. Although motor training paradigms, similar to the one employed in the present study, have been shown to induce LTP-like changes in cortical activity [Ziemann et al., 2004], it would be fair to consider this hypothesis speculative if based on fMRI alone. In the present work, however, it is supported be two additional lines of evidence: changes in TMS-evoked responses, and changes in cortical thickness.

### TMS

TMS provides an indirect way to assess LTP-like changes in cortical excitability. Since TMS activates motor cortical output neurons trans-synaptically, if synaptic efficacy is increased, this should lead to an increase in the amplitude of the MEP at a given stimulus intensity [Di Lazzaro and Ziemann, 2013]. Here, we showed that MEP amplitudes were larger following training (Figure 4). Notably, there was no significant change in the area that could evoke an MEP in the APB. This suggests that the changes indexed by TMS were predominantly driven by changes in neural networks that already played a role in contraction of the APB.

In order to consider TMS and MRI evidence together, it is important that these are viewed with respect to one another, to ascertain that regions of measurement (or signal change) for these modalities have reasonable anatomical overlap. In the present study, the areas of cortex that were activated by the trained sequence (in the trained hemisphere) during fMRI scans were very similar, though not identical, to areas eliciting MEPs during TMS mapping (Figure 5). TMS and fMRI map markedly different aspects of motor control, and so perfect overlap between them should not be expected: fMRI here contrasted BOLD responses to sequential movement of the fingers and thumb versus rest, while TMS targeted neurons functionally relevant to activating the APB muscle.

Keeping in mind that there was no increase in the RMT, the TMS findings suggest that at least one of three processes have taken place: enhanced trans-synaptic transmission (e.g., through LTP), increased neurite density, and/or improved conduction of the corticospinal tract. Both the first and second of these possibilities support our earlier hypothesis introduced in the context of the fMRI results. The third possibility, regarding changes in white matter, is investigated in detail in our follow-up publication [Reid et al.,].

### Cortical Thickness

Increased cortical thickness was observed after motor training within the right SMA, right middle-frontal gyrus, as well as right pre-central gyrus and right post-central gyrus. Although cortical thickness changes did not reach our stricter FWE-corrected threshold, the clusters of significant voxels seen were sizeable (22 - 81 voxels @ p<0.001) and striking in their location. The location of clusters in the pre-and post-central gyri were consistent with our fMRI maps, our TMS maps (Figure 6), and with areas responsible for sensorimotor representation of the fingers elucidated through electrocorticographic and brain stimulation techniques [Penfield and Boldrey, 1937; Wahnoun et al., 2015]. The neighbouring clusters seen in the right SMA were also within the TMS and fMRI maps from our study, and consistent with previous findings that this area plays a role in motor learning [Taubert et al., 2010]. That these changes appeared near the peaks of fMRI and TMS maps adds further credibility to the suggestion that changes in fMRI and TMS patterns were a reflection of altered neurite organisation, density, and/or connection strength in regions that, at baseline, were already predominantly responsible for execution of the (to-be) learned sequence, rather than altered responsibility of surrounding areas.

The final location in which we saw changes in cortical thickness was the dlPFC, which is known to play an important role in error-correction of motor output (along with the caudate nucleus) [Chevrier et al., 2007; Kübler et al., 2006]. Structural changes in the dlPFC were toward the edge of the TMS response map, and not in an area of fMRI change. The role of the dlPFC is to *modulate* motor responses, not generate them directly, and so it should not be surprising that only small TMS responses were recorded. Furthermore, we do not consider cortical thickness change here to be at odds with fMRI findings because the fMRI analysis was optimised for detection of motor output and conducted at around half the speed participants were capable of at baseline; it did not contrast very-challenging-versus-relaxed motor performance, as would be optimal to highlight an area that plays a role in error detection. In fact, we believe this apparent discrepancy highlights the usefulness of multimodality imaging in measuring neuroplasticity to provide a more accurate overview of changes that take place during learning [Reid et al., 2016]. The dlPFC and related areas were further investigated with diffusion imaging; these analyses are described in a separate publication [Reid et al.,].

### Conclusion

We have shown that four weeks of motor training can invoke robust and reliable changes in behaviour, cortical thickness, TMS-evoked motor maps, and fMRI activation in motor regions of the brain. Taken together with our diffusion MRI findings [Reid et al.,], the work presented here provides the most cohesive and comprehensive longitudinal study of motor plasticity in healthy adult humans to date. All three brain measures suggested that motor learning was driven by LTP-like plasticity that was relatively widespread across the sensorimotor system – even when a participant is trained on a simple task solely on the basis of proprioceptive feedback.

## Acknowledgements

This work was supported by a Training Fellowship from the National Health and Medical Research Council of Australia (APP1012153) awarded to MVS. JBM was supported by an Australian Research Council Laureate Fellowship (FL110100103). The funders had no role in study design, data collection and analysis, decision to publish or preparation of the manuscript. The content is solely the responsibility of the authors and does not necessarily represent the official views of the funding bodies. The authors declare no competing financial interests. We would like to thank Dr. Amanda Robinson, Dr. David Lloyd, and Dr. Daniel Stjepanovic for technical assistance.

## Supplementary Material

**Supplementary Figure 1.**
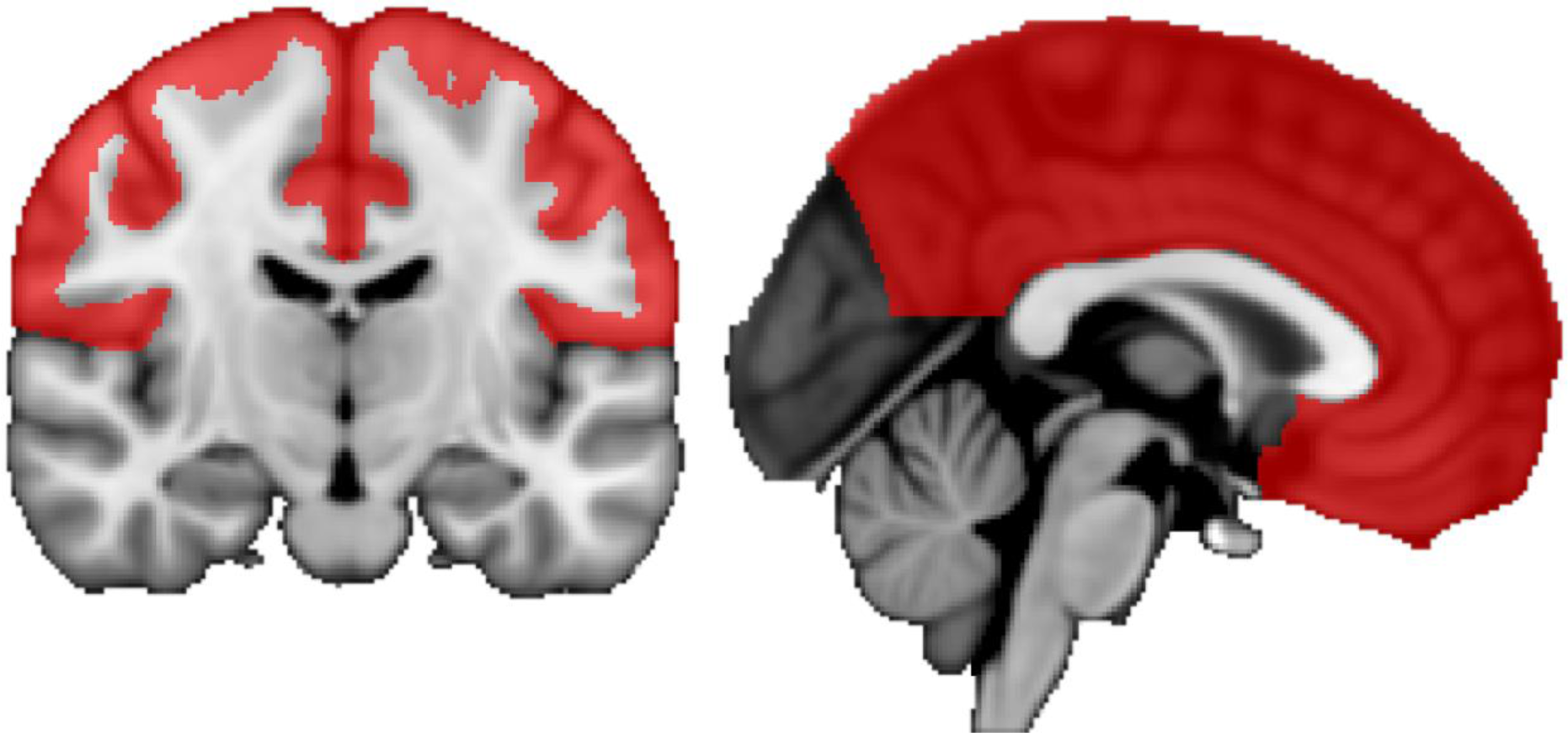
Region of interest for cortical thickness analysis (red) overlaid on the FSL MNI template used to coregister participants. To correct segmentation errors, brain masks for single-subject templates were manually edited in every sagittal, axial, and coronal slice prior to warping to MNI space. Final segmentation acceptability was based predominantly on the frontal and parietal lobes (red). Voxel-based morphology statistics were solely calculated for these regions.

**Supplementary Figure 2.**
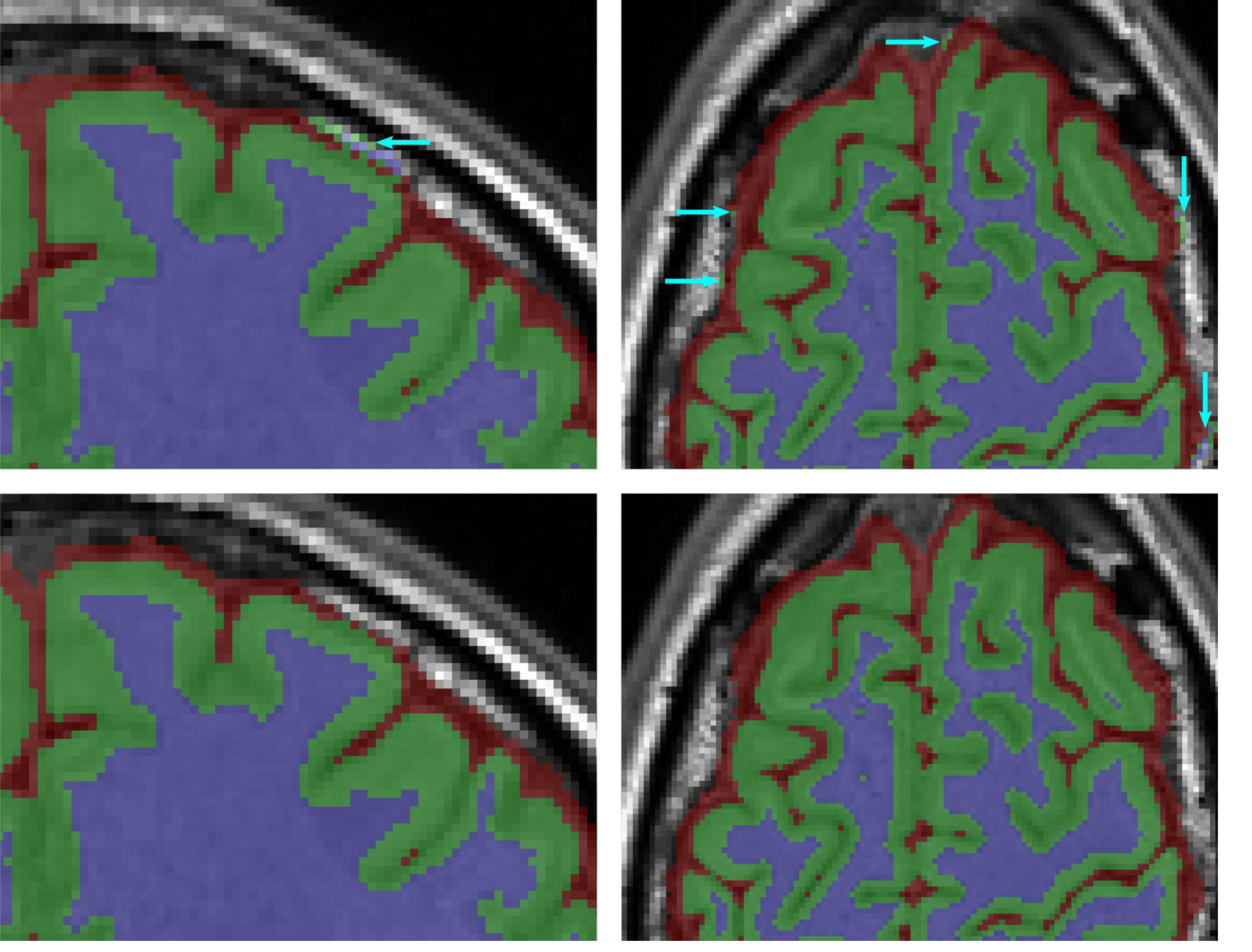
Coronal (left column) and Axial (right column) views of final hard segmentations achieved for a single-subject template, as calculated by ANTs. Blue, green, and red indicate white matter, grey matter, CSF, respectively. **Top Row:** segmentation performed without any form of manual correction. Arrows indicate locations where dura has been categorised as grey and/or white matter. The existence of any of these errors would deem the segmentation unacceptable in the present study. Small (<5 voxel) clusters of grey matter labelling of the dura in the temporal and occipital lobes were generally corrected but did not automatically deem an entire segmentation as unacceptable if the only error present. **Bottom Row:** segmentation of the same template using repeated brain mask correction to correct segmentation errors. The displayed images are representative of the typical error-correction performed, and final segmentation quality obtained, in the present study. Left of images is subject’s left. Top of axial images is anterior. Top of coronal images is superior.

## References

Amunts K, Schlaug G, Jäncke L, Steinmetz H, Schleicher A, Dabringhaus A, Zilles K (1997): Motor cortex and hand motor skills: structural compliance in the human brain. Hum Brain Mapp 5:206–15. http://www.ncbi.nlm.nih.gov/pubmed/20408216.

Avants BB, Epstein CL, Grossman M, Gee JC (2008): Symmetric diffeomorphic image registration with cross-correlation: evaluating automated labeling of elderly and neurodegenerative brain. Med Image Anal 12:26–41. http://www.ncbi.nlm.nih.gov/pubmed/17659998.

Barron HC, Vogels TP, Emir UE, Makin TR, O’Shea J, Clare S, Jbabdi S, Dolan RJ, Behrens TEJ (2016): Unmasking Latent Inhibitory Connections in Human Cortex to Reveal Dormant Cortical Memories. Neuron 90:191–203. http://dx.doi.org/10.1016/j.neuron.2016.02.031.

Bezzola L, Mérillat S, Gaser C, Jäncke L (2011): Training-induced neural plasticity in golf novices. J Neurosci 31:12444–8. http://www.jneurosci.org/cgi/doi/10.1523/JNEUROSCI.1996-11.2011.

Bliss T V, Collingridge GL (1993): A synaptic model of memory: long-term potentiation in the hippocampus. Nature 361:31–9. http://www.ncbi.nlm.nih.gov/pubmed/15190253.

Chang Y (2014): Reorganization and plastic changes of the human brain associated with skill learning and expertise. Front Hum Neurosci 8:35. http://journal.frontiersin.org/Article/10.3389/fnhum.2014.00035/abstract.

Chevrier AD, Noseworthy MD, Schachar R (2007): Dissociation of response inhibition and performance monitoring in the stop signal task using event-related fMRI. Hum Brain Mapp 28:1347–1358.

Costa RM, Cohen D, Nicolelis MAL (2004): Differential corticostriatal plasticity during fast and slow motor skill learning in mice. Curr Biol 14:1124–34. http://linkinghub.elsevier.com/retrieve/pii/S0960982204004658.

Draganski B, Gaser C, Busch V, Schuierer G, Bogdahn U, May A (2004): Neuroplasticity: changes in grey matter induced by training. Nature 427:311–2. http://www.ncbi.nlm.nih.gov/pubmed/14737157.

Floyer-Lea a, Matthews PM (2005): Distinguishable brain activation networks for short-and long-term motor skill learning. J Neurophysiol 94:512–8. http://www.ncbi.nlm.nih.gov/pubmed/15716371.

Hallett M (2000): Transcranial magnetic stimulation and the human brain. Nature 406:147–50. http://www.nature.com/doifinder/10.1038/35018000.

Hoy RR, Nolen TG, Casaday GC (1985): Dendritic sprouting and compensatory synaptogenesis in an identified interneuron follow auditory deprivation in a cricket. Proc Natl Acad Sci U S A 82:7772–6. http://www.pubmedcentral.nih.gov/Articlerender.fcgi?artid=391416&tool=pmcentrez&rendertype=abstract.

Karni A, Meyer G, Jezzard P, Adams MM, Turner R, Ungerleider LG (1995): Functional MRI evidence for adult motor cortex plasticity during motor skill learning. Nature 377:155–8. http://lbcnimh.nih.gov/Ungerleider/Publications/Karni_et_al_Nature_1995.pdf.

Kong NW, Gibb WR, Tate MC (2016): Neuroplasticity: Insights from Patients Harboring Gliomas. Neural Plast 2016:2365063. http://www.hindawi.com/journals/np/2016/2365063/.

Kübler A, Dixon V, Garavan H (2006): Automaticity and reestablishment of executive control-an fMRI study. J Cogn Neurosci 18:1331–42. http://www.ncbi.nlm.nih.gov/pubmed/16859418.

Lancaster JL, Tordesillas-Gutiérrez D, Martinez M, Salinas F, Evans A, Zilles K, Mazziotta JC, Fox PT (2007): Bias between MNI and Talairach coordinates analyzed using the ICBM-152 brain template. Hum Brain Mapp 28:1194–205. http://www.ncbi.nlm.nih.gov/pubmed/17266101.

Di Lazzaro V, Ziemann U (2013): The contribution of transcranial magnetic stimulation in the functional evaluation of microcircuits in human motor cortex. Front Neural Circuits 7:18. http://www.ncbi.nlm.nih.gov/pubmed/23407686.

Logothetis NK (2008): What we can do and what we cannot do with fMRI. Nature 453:869–78. http://www.ncbi.nlm.nih.gov/pubmed/18548064.

Luders E, Narr KL, Thompson PM, Rex DE, Woods RP, DeLuca H, Jancke L, Toga AW (2006): Gender effects on cortical thickness and the influence of scaling. Hum Brain Mapp 27:314–324.

Matsuzaki M, Honkura N, Ellis-Davies GCR, Kasai H (2004): Structural basis of long-term potentiation in single dendritic spines. Nature 429:761–6. http://www.ncbi.nlm.nih.gov/pubmed/15190253.

Milton J, Solodkin A, Hlustík P, Small SL (2007): The mind of expert motor performance is cool and focused. Neuroimage 35:804–13. http://www.ncbi.nlm.nih.gov/pubmed/17317223.

Nudo RJ, Wise BM, SiFuentes F, Milliken GW (1996): Neural substrates for the effects of rehabilitative training on motor recovery after ischemic infarct. Science 272:1791–4. http://www.ncbi.nlm.nih.gov/pubmed/8650578.

Pascual-Leone A, Amedi A, Fregni F, Merabet LB (2005): The plastic human brain cortex. Annu Rev Neurosci 28:377–401.

Pearce a J, Thickbroom GW, Byrnes ML, Mastaglia FL (2000): Functional reorganisation of the corticomotor projection to the hand in skilled racquet players. Exp brain Res 130:238–43. http://www.ncbi.nlm.nih.gov/pubmed/10672477.

Penfield W, Boldrey E (1937): Somatic Motor and Sensory Representation in the Cerebral Cortex of Man as Studies by Electrical Stiumlation. Brain 60:389–443. http://brain.oxfordjournals.org/cgi/doi/10.1093/brain/60.4.389.

Reid LB, Boyd RN, Cunnington R, Rose SE (2016): Interpreting Intervention Induced Neuroplasticity with fMRI: The Case for Multimodal Imaging Strategies. Neural Plast 2016:1–13. http://www.hindawi.com/journals/np/2016/2643491/.

Reid LB, Rose SE, Boyd RN (2015): Rehabilitation and neuroplasticity in children with unilateral cerebral palsy. Nat Rev Neurol 11:390–400. http://www.nature.com/doifinder/10.1038/nrneurol.2015.97.

Reid LB, Sale M V, Cunnington R, Mattingley JB, Rose SE Structural and functional brain changes following four weeks of unimanual motor training: evidence from fMRI-guided diffusion MRI tractography. Hum Brain Mapp VOLUME:PAGES.

Rioult-Pedotti M-SS, Friedman D, Donoghue JP (2000): Learning-induced LTP in neocortex. Science (80-) 290:533–6. http://www.sciencemag.org/cgi/doi/10.1126/science.290.5491.533.

Roth Y, Amir A, Levkovitz Y, Zangen A (2007): Three-dimensional distribution of the electric field induced in the brain by transcranial magnetic stimulation using figure-8 and deep H-coils. J Clin Neurophysiol 24:31–8. http://www.ncbi.nlm.nih.gov/pubmed/17277575.

Sale M V., Ridding MC, Nordstrom MA (2008): Cortisol inhibits neuroplasticity induction in human motor cortex. J Neurosci 28:8285–93. http://www.jneurosci.org/cgi/doi/10.1523/JNEUROSCI.1963-08.2008.

Sanes JN, Donoghue JP (2000): Plasticity and primary motor cortex. Annu Rev Neurosci 23:393–415. http://www.ncbi.nlm.nih.gov/pubmed/10845069.

Schabrun SM, Stinear CM, Byblow WD, Ridding MC (2009): Normalizing Motor Cortex Representations in Focal Hand Dystonia. JOUR. Cereb Cortex 19:1968–1977. http://cercor.oxfordjournals.org/content/19/9/1968.abstract.

Scholz J, Klein MC, Behrens TEJ, Johansen-Berg H (2009): Training induces changes in white-matter architecture. Nat Neurosci 12:1370–1371. http://www.pubmedcentral.nih.gov/Articlerender.fcgi?artid=2770457&tool=pmcentrez&rendertype=abstract.

Stefan K, Kunesch E, Cohen LG, Benecke R, Classen J (2000): Induction of plasticity in the human motor cortex by paired associative stimulation. Brain 123 Pt 3:572–584.

Taubert M, Draganski B, Anwander A, Müller K, Horstmann A, Villringer A, Ragert P (2010): Dynamic properties of human brain structure: learning-related changes in cortical areas and associated fiber connections. J Neurosci 30:11670–7. http://www.ncbi.nlm.nih.gov/entrez/query.fcgi?cmd=Retrieve&db=PubMed&dopt=Citation&list_uids=20810887.

Thomas AG, Marrett S, Saad ZS, Ruff DA, Martin A, Bandettini PA (2009): Functional but not structural changes associated with learning: An exploration of longitudinal Voxel-Based Morphometry (VBM). Neuroimage 48:117–125. http://dx.doi.org/10.1016/j.neuroimage.2009.05.097.

Wahnoun R, Benson M, Helms-Tillery S, Adelson PD (2015): Delineation of somatosensory finger areas using vibrotactile stimulation, an ECoG study. Brain Behav 5:e00369. http://www.ncbi.nlm.nih.gov/pubmed/26516605.

Xiong J, Ma L, Wang B, Narayana S, Duff EP, Egan GF, Fox PT (2009): Long-term motor training induced changes in regional cerebral blood flow in both task and resting states. Neuroimage 45:75–82. http://www.pubmedcentral.nih.gov/Articlerender.fcgi?artid=2672588&tool=pmcentrez&rendertype=abstract.

Xu T, Yu X, Perlik AJ, Tobin WF, Zweig J a, Tennant K, Jones T, Zuo Y (2009): Rapid formation and selective stabilization of synapses for enduring motor memories. Nature 462:915–9. http://www.pubmedcentral.nih.gov/Articlerender.fcgi?artid=2844762&tool=pmcentrez&rendertype=abstract.

Yushkevich P a., Piven J, Hazlett HC, Smith RG, Ho S, Gee JC, Gerig G (2006): User-guided 3D active contour segmentation of anatomical structures: significantly improved efficiency and reliability. Neuroimage 31:1116–28. http://www.ncbi.nlm.nih.gov/pubmed/16545965.

Ziemann U, Ilić T V, Pauli C, Meintzschel F, Ruge D (2004): Learning modifies subsequent induction of long-term potentiation-like and long-term depression-like plasticity in human motor cortex. J Neurosci 24:1666–72. http://www.jneurosci.org/cgi/doi/10.1523/JNEUROSCI.5016-03.2004.

